# Benchmarking Oxford Nanopore read assemblers for high-quality molluscan genomes

**DOI:** 10.1101/2020.12.31.424979

**Authors:** Jin Sun, Runsheng Li, Chong Chen, Julia D. Sigwart, Kevin M. Kocot

**Affiliations:** Institute of Evolution & Marine Biodiversity, Ocean University of China, Qingdao 266003, China; Department of Infectious Diseases and Public Health, Jockey Club College of Veterinary Medicine and Life Sciences, City University of Hong Kong, Kowloon, Hong Kong, China; X-STAR, Japan Agency for Marine-Earth Science and Technology (JAMSTEC), 2-15 Natsushima-cho, Yokosuka, Kanagawa Prefecture 237-0061, Japan; Senckenberg Museum, Frankfurt, Germany; Queen’s University Belfast, Marine Laboratory, Portaferry, N Ireland; Department of Biological Sciences and Alabama Museum of Natural History, University of Alabama, Tuscaloosa, Alabama, 35487, USA

**Keywords:** Molluscan genomes, assembly, Oxford Nanopore Technology, scaly-foot snail, Mytilus, phylogeny

## Abstract

Choosing the optimum assembly approach is essential to achieving a high-quality genome assembly suitable for comparative and evolutionary genomic investigations. Significant recent progress in long-read sequencing technologies such as PacBio and Oxford Nanopore Technologies (ONT) also brought about a large variety of assemblers. Although these have been extensively tested on model species such as *Homo sapiens* and *Drosophila melanogaster*, such benchmarking has not been done in Mollusca which lacks widely adopted model species. Molluscan genomes are notoriously rich in repeats and are often highly heterozygous, making their assembly challenging. Here, we benchmarked 10 assemblers based on ONT raw reads from two published molluscan genomes of differing properties, the gastropod *Chrysomallon squamiferum* (356.6Mb, 1.59% heterozygosity) and the bivalve *Mytilus coruscus* (1593Mb, 1.94% heterozygosity). By optimising the assembly pipeline, we greatly improved both genomes from previously published versions. Our results suggested that 40-50X of ONT reads are sufficient for high-quality genomes, with Flye being the recommended assembler for compact and less heterozygous genomes exemplified by *C. squamiferum*, while NextDenovo excelled for more repetitive and heterozygous molluscan genomes exemplified by *M. coruscus*. A phylogenomic analysis utilising the two updated genomes with other 32 published high-quality lophotrochozoan genomes resulted in maximum support across all nodes, and we show that improved genome quality also leads to more complete matrices for phylogenomic inferences. Our benchmarking will ensure the efficiency in future assemblies for molluscs and perhaps also other marine phyla with few genomes available.

## Introduction

Sequencing the whole genome of an organism can be highly beneficial to understanding its biology and evolution. High-quality genome assemblies allows researchers to begin deciphering the causal connection between genotype and phenotype, explore the molecular control of traits through gene expression and regulation, shed light on the species’ evolutionary process at the genomic scale, and perform comprehensive and robust phylogenomic analyses to infer evolutionary relationships [1]. Mollusca is the second largest animal phylum, and their habitats cover a dramatic range of ecosystems worldwide, from high mountains to the deep sea, from lush rainforests to arid deserts, and from freshwater streams to coral reefs. Many molluscs provide important resources for humans, being the largest aquaculture resources second only to teleost fishes [2]; a number of disease-causing human parasites also use molluscs as an intermediate host, such as blood flukes in the genus *Schistosoma* which causes schistosomiasis [3]. With members as different as the colossal squid and microscopic worms living between grains of sand, it is also the most morphologically disparate animal phylum [4], and the understanding of their biology and evolution provides clues to answer fundamental questions on genotypic vs phenotypic adaptation, origins of evolutionary novelties, and macroevolutionary processes.

Currently, the number of published molluscan genomes is relatively small compared to other major phyla such as Arthropoda and Chordata. At the time of writing (September 2020), there are 666 Arthropoda and 1372 Chordata genomes deposited on NCBI Genome database; only 29 high-quality Mollusca genomes (i.e. BUSCO score over 80%) have been published (Supplementary Table S1), representing only approximately <0.05% of molluscan species (considering the number of described species in Mollusca is ~65,000). Among these, only 9 are chromosome-scale. In addition, the sequenced taxa are heavily biased to the conchiferan classes of Bivalvia, Gastropoda, and, to a lesser extent, Cephalopoda, with no published high-quality genomes in other classes including the aculiferans (Polyplacophora, Solenogastres, and Caudofoveata), which are key to understanding molluscan evolution. The small number of molluscan genomes sequenced and the taxonomic bias can be attributed to a few reasons: 1) some groups, such as monoplacophorans, are rare and not easy to collect [5]; 2) extraction of adequate, high-quality genomic DNA from molluscs can be very difficult; 3) molluscan genomes tend to be very heterozygous and repetitive, which significantly hinders the assembly of reads into high-quality genomes.

Along with the rapid progress of sequencing technologies in recent years, especially so-called third-generation long read sequencing technologies [PacBio and Oxford Nanopore Technologies (ONT)], a large variety of assemblers have been developed for long-read based genome assembly. Although the effectiveness of different assemblers has been extensively tested on model species such as *Homo sapiens* and *Drosophila melanogaster*, there are currently no widely adopted molluscan model species. As such, the efficacy of different genomic assemblers for *de novo* molluscan genome assembly has not been evaluated systematically. Considering the distinct genomic features between molluscan genomes sequenced to date and model species in other phyla, the assembly strategies and assemblers effective in those model species are unlikely to be equally effective for molluscan genomes. Since running and testing different assemblers is computationally intensive and time consuming, it is beneficial to carry out a systematic benchmarking of different assembly strategies for molluscs to identify the best strategy for assembling future molluscan genomes.

Here, we selected two species from different molluscan classes with existing genomes that were previously assembled from ONT reads as the models to benchmark different assembly strategies. Namely, we focused on the Scaly-foot Snail *Chrysomallon squamiferum* (Gastropoda) and the Hard-shelled Mussel *Mytilus coruscus* (Bivalvia). We selected ONT because 1) the low cost of the MinION instrument makes this technology more accessible to researchers studying molluscs, 2) currently more tools have been specifically designed for ONT data, such as NECAT [6], NextDenovo (https://github.com/Nextomics/NextDenovo), and Shasta [7], and 3) the potential to apply the ultra-long DNA sequencing capability of ONT. The published Scaly-foot Snail genome is relatively compact (444.4 Mb) while the *M. coruscus* genome is larger (1.90 Gb); both genomes are very heterozygous, with the heterozygosity of *C. squamiferum* being 1.38% and *M. coruscus* being even higher at 1.64% [8, 9]. We recorded genome contiguity, completeness, as well as misassemblies resulting from up to 10 assemblers. Using these two re-assembled and improved genomes, we performed an updated phylogenomic analysis with all publicly available, high-quality molluscan genomes and explored the gene families specifically expanded in particular lineages within Mollusca, exemplifying the utility of high-quality genomes in understanding the evolution and biology of molluscs.

## Methods

Raw ONT reads in the *fast5* format from the published Scaly-foot Snail genome sequencing project (PRJNA523462) [8] were re-basecalled using Guppy version 3.6.0 with the high-accuracy (HAC) mode on a GeForce® RTX 1080 Ti (NVIDIA) GPU. The Illumina and ONT reads from the *Mytilus coruscus* genome sequencing project were downloaded from the NCBI SRA database (ERR3415816 and ERR3431204)[9]. Illumina reads were cleaned to remove bacterial contamination using Kraken2 [10], and the genome size and heterozygosity was calculated by Jellyfish v2.3.0 with the k-mer size of 17, 19, and 21 and GenomeScope 2.0 [11].

The following assemblers were used for the benchmarking, including the long-read only assemblers (Canu [12], Flye [13], Wtdbg2 [14], Miniasm [15], NextDenovo (https://github.com/Nextomics/NextDenovo), NECAT [6], Raven [16], and Shasta [7]) and hybrid assemblers (MaSuRCA [17] and QuickMerge [18]). Canu was not tested on the *M. coruscus* genome due to the extremely intensive computing time required for this large genome. Previous analyses have suggested using corrected ONT reads could improve the genome assembly [19]. To check the effect of this has on the assemblies, the ONT reads that were corrected and / or trimmed by Canu and NECAT were also tested. To check whether including the shorter ONT reads could affect the assembly. The ONT reads were also sub-sampled with different cut-off lengths (see Table 1 for the lengths used). CPU hour was calculated in the Slurm workload manager system by recording the program start and end time point. However, since the hardware configuration in each node varied, the CPU hour presented is only an indicator of the relative trend among different assemblers.

**Table 1.**
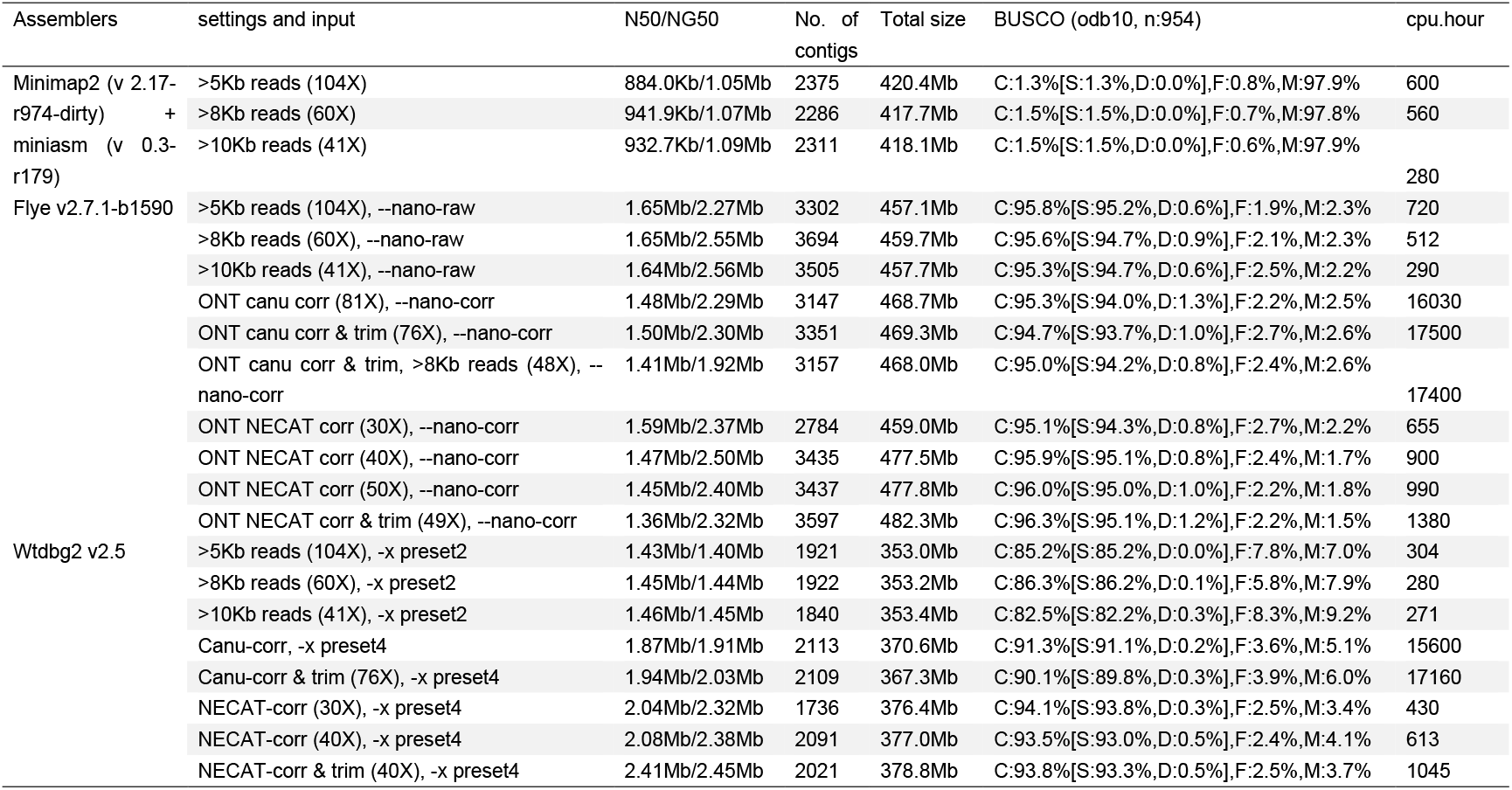
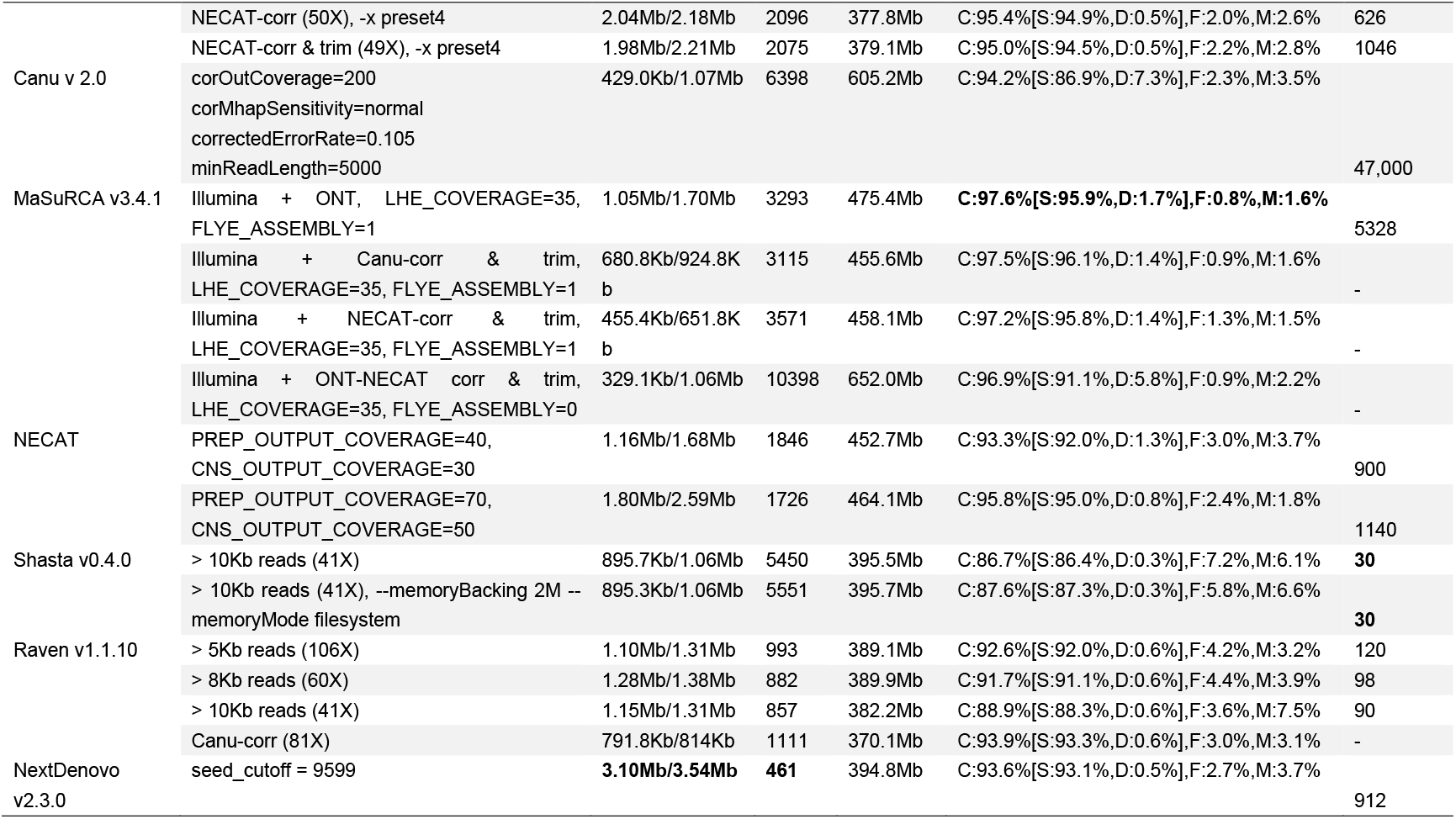
Assembly results for the Scaly-foot snail genome. The best performance was highlighted in bold. Corr, corrected reads, trim, corrected and trimmed reads. NG50 is the N50 value after normalization using the predicted genome size.

The assembled contigs were polished at least three times with Flye, and heterozygous contigs were removed with the purge_dup pipeline [20]. The resultant genomes were polished twice using Pilon version 1.23 [21] with Illumina reads. The genome completeness of each assembly was thoroughly monitored at each step using BUSCO v4.0.6 with odb10 metazoan dataset [22]. The genome quality of the Scaly-foot Snail assemblies were assessed by QUAST version 5.0.2 [23] comparing against the formerly published version of the genome as a reference [8]. QUAST calculates genome assembly characteristics such as N50 and total size, but also assesses mis-assemblies with minimap2. The detailed commands and settings used for all analyses can be found in the Supplementary Information.

Repeat content was initially predicted with RepeatModeler version 2.0.1. Genomes were hard-masked using RepeatMasker v4.1.0 with a species-specific repeat library generated by RepeatModeler and all the known repeat content in the RepeatMasker repeat database. Augustus version 3.3.3, a *de novo* gene predictor, was trained using Braker version 2.1.5 with the hard-masked genome and the transcriptome assemblies. Genome annotation was then performed using Maker v3.01.03 with the trained Augustus *ab initio* predictor plus each species’ transcriptome assembly and molluscan protein sequences downloaded from the NCBI protein database (July 2020). Each gene was annotated by InterProScan-5.36-75.0.

To identify putative orthologous sequences shared among taxa, we used OrthoFinder 2.4.0 [24] with an inflation parameter of 2.1. Working from the *.fasta* files generated in the “Orthogroup_Sequences” directory, we removed sequences shorter than 100 amino acids and removed sequences that were identical where they overlapped, keeping the longest non-redundant sequence. We then retained only those *.fasta* files sampled for at least four taxa, and aligned them using MAFFT 7.310 [25] with the following options: --auto, --localpair, and --maxiterate 1000. We then removed putatively mistranslated regions with HmmCleaner [26] with the --specificity option. We deleted sequences that did not overlap with any other sequences by at least 20 amino acids, starting with the shortest sequence not meeting this overlap criterion. Then, we trimmed the alignments to remove ambiguously aligned and ‘noisy’ regions with BMGE v. 1.12.2 [27] and constructed “approximately maximum likelihood” trees for each alignment with FastTree 2 [28] using the -slow and -gamma options. In order to identify strictly orthologous sequences among taxa, we used PhyloPyPruner 0.9.5 (https://pypi.org/project/phylopypruner) with the following options: --min-support 0.9 --mask pdist --trim-lb 3 --trim-divergent 0.75 --min-pdist 0.01 --prune LS. Only alignments sampled for at least 75% of the taxa (i.e., 26 taxa) were retained for the final analyses. Datasets with genes sampled for at least four taxa and at least 50% of the taxa were also generated (available in the supplementary materials). In order to check if higher-quality genome assemblies lead to the recovery of more orthogroups, a regression analysis was carried out in SPSS 16.0 between BUCSO scores and the orthologue gene occupancy.

Phylogenetic analyses were conducted on the partitioned supermatrix produced by PhyloPyPruner using IQ-Tree 2 [29] with the best-fitting model of amino acid evolution for each partition (-m MFP). Topological support was assessed with 1,000 rapid bootstraps. MCMCTree version 4.8a was used to calibrate the time constraints. The “root-age” was set as 590Ma [30]. The following time constraints were applied: a soft minimum bound of 245 Ma for the first appearance of Ostreoidea [31]; a hard minimum bound of 465 Ma for the first appearance of Pteriomorpha [32]; a soft minimum bound of 125 Ma for the first appearance of Mactroidea [31]; a soft constraint of 520.5 Ma and 530 Ma for the origin of Bivalvia [31]; a hard upper bound of 150 Ma for the split of *Lanistes nyassanus* (representing the old world ampullariids) and the new world ampullariids [33]; a hard lower bound of 130 Ma for the first appearance of both the Stylommatophora and Hygrophila [34]; a hard lower bound of 168.6 Ma and a soft upper bound of 473.4 Ma for the split of *Aplysia* and *Biomphalaria* [35]; a hard minimum 390 Ma bound for the split of Caenogastropoda and Heterobranchia [36]; a hard lower bound of 470.2 Ma and a soft upper bound of 531.5 Ma for the first appearance of Gastropoda [35]; a hard lower bound of 532 Ma and a soft upper bound of 549 Ma for the first appearance of molluscs [37]; a hard lower bound of 550.25 Ma and a hard upper bound of 636.1 Ma for the origin of Lophotrochozoa [37]. The model of LG+I+G, which was the best-fitting model for the vast majority of the single-gene partitions, was applied with the burn-in set to 10 million and the sampling frequency set to 1000.

## Results and Discussions

### (a) Scaly-foot Snail genome assembly

The Illumina sequencing reads used for the previously published genome assembly of *Chrysomallon squamiferum* were found to contain some endosymbiont contamination, which likely led to the overestimation of the genome size (444.4 Mb). With cleaned Illumina reads, the genome size was predicted to be 356.6 Mb, and the heterozygosity was estimated to be 1.59%. Although k-mer count methods for genome size estimation may still be biased by the high heterozygosity, we used this genome size in the downstream analyses.

The high-accuracy mode of Guppy 3.6.0 significantly improved the ONT read accuracy, with the base quality score being improved from 13.2±1.5 when not using the high-accuracy mode to 16.9±3.5 in the case of the Scaly-foot Snail genome. Different genome assemblies were tested on the newly basecalled ONT reads.

The assembly results from different assemblers using different settings and filters are shown in Table 1. In the case of the minimap2+miniasm assembly, the best assembly resulted from >10Kb or longest 41X filtering ONT reads, suggesting the inclusion of shorter reads may actually reduce the contiguity of the assembly. A similar observation was also reported in a study assembling the *Caenorhabditis elegans* genome with ONT reads [38]. However, due to Miniasm lacking a sequence consensus step, the base accuracy in the assembly can only be as good as the input reads, leading to a very poor BUSCO score (1.5%) (Table 1, Supplementary Table S2). Similarly, the best assembly using Flye resulted from filtering the raw ONT reads keeping only those longer than 10Kb; including shorter raw reads also appeared to reduce the contiguity of the assembly. Output assembly quality from Flye also seemed to be independent from whether the reads were corrected and/or trimmed or not, though the assembly with the longest 50X NECAT-corrected ONT reads had the longest contig (9.84Mb) among the assemblies done with Flye (Supplementary Table S2). These results indicate that Flye is indeed mainly optimised for raw ONT reads, as suggested by its authors [13]. For Wtdbg2, increasing the length filter also increased the assembly contiguity. The >10kb filter for raw ONT reads resulted in a higher-quality assembly than the >8kb for and >5Kb filter. Both Canu-corrected and NECAT-corrected ONT reads dramatically increased the NG50, and the trimming on the corrected reads to remove suspicious reads (e.g. adapters) was also effective in increasing the assembly quality. The assembly using the longest 49X NECAT-corrected and trimmed reads had the longest contig (10.64Mb), and the assembly using the longest 40X NECAT-corrected and trimmed reads had the best NG50 value (2.44Mb).

Among the MaSuRCA assemblies, the highest quality assembly was from combining Illumina reads together with the longest 35X raw ONT reads; assemblies with Illumina reads and corrected/trimmed ONT reads did not increase the NG50. The MaSuRCA assembly had the highest BUSCO score (97.6%) at this point. This suggests a high base accuracy, as the mega-reads generated by MaSuRCA effectively combined both the more accurate Illumina reads and the longer ONT reads [17]. With the NECAT assembler, using the longest 50X raw ONT reads improved the assembly compared to when using top 30x, suggesting longest 50x reads may be the better input for NECAT. For Shasta assemblies, using the default settings and the settings of “-memoryMode filesystem and -memoryBacking 2M” resulted in assemblies of similar qualities. This result is different from what other authors reported in other genome assemblies of model species such as human and *Drosophila* [7] where the latter setting resulted in much improved assemblies, suggesting that the latter settings may not be beneficial for highly heterozygous genomes like those of Mollusca. The final assembler tested was Raven, an updated version of Ra [16]. The best assembly resulted from using either >8Kb or longest 60X filter ONT reads, running the assembler with Canu-corrected reads actually resulted in lower quality genome assemblies (Table 1).

With regards to the computing time (Table 1), Shasta was the fastest assembler, followed by Raven, minimap2+miniasmand, and Wtdbg2. Canu was the most computationally intensive, followed by MaSuRCA, then NECAT or NextDenovo. Canu also took more CPU hours to correct the reads than NECAT.

As different assemblers may include different (or lacking) consensus steps, and some assemblers may merge the heterozygous contigs (or ‘bubble’ in the assembly), the best genome assembly from each assembler (as judged by NG50, the N50 value after normalization using the predicted genome size) was polished with ONT reads using Flye. Removal of heterozygous contigs by purge_dups was carried out when the coverage histogram of the mapped ONT reads exhibited a heterozygous peak, which was often necessary except for Wtdbg2, Raven, and Shasta, suggesting that these three assemblers were able to actively merge heterozygous contigs. Furthermore, this step also helped to increase the BUSCO score by decreasing the duplicated BUSCOs (Supplementary Table S3). Among the post-polishing assemblies, the Flye assembly had the highest BUSCO score (C:97.8% [D:0.6%]). This is in line with the published finding that Flye is capable of assembling some genomic regions that may be missing in assemblies produced by other assemblers by better resolving the repetitive regions [13]. Following Flye, the next most complete assembly was that from minimap2+Miniasm (C:97.8% [D:0.7%]), and then NECAT and Shasta (both C:97.6%[D:0.4%]).

In terms of genome contiguity, the assembly from QuickMerge (merging the Flye version and MaSuRCA version) exhibited the highest NG50 (4.00Mb), followed by NextDenovo (3.40Mb), Flye (2.55Mb) and NECAT (2.50Mb).

Analysis of misassembly and mismatch with QUAST revealed that NextDenovo has the least number of misassemblies, followed by QuickMerge. For the number of mismatches per 100 kb, Raven was the best performer followed by NextDenovo; for number of indels per 100 kb, MaSuRCA performed the best, followed by Shasta. Nevertheless, for genomes resulting from these 10 assemblers, the amount of misassembly, mismatches, and indels were not very different, particularly the latter two parameters (number of mismatches and number of indels per 100kb), indicating that they were similar in performance.

We selected the polished Flye assembly for the downstream analysis due to this assembly exhibiting the highest BUSCO score and also the largest size among the assemblies. The Hi-C library from the original published assembly [8] was used to further scaffold the contigs in the Flye assembly, resulting in a final assembly including 15 pseudo-chromosomal scaffolds plus 492 contigs. Annotating the genome suggested a dramatic improvement compared to the original published assembly in the number of gene models (21,469 vs 16,917), and the BUSCO score of the predicted genes increased from 87.5% to 94.1%. This assembly is one of the most complete genome assemblies in Mollusca to date.

### *(b)* Mytilus coruscus *genome assembly*

The genome size of *M. coruscus* predicted with GenomeScope 2.0 was 1,593Mb, smaller than the size predicted by an earlier study (1.85Gb), and heterozygosity was estimated to be 1.94% [9]. A total of 158.9 Gb of ONT reads (99.8 X) with the base quality score of 13.1±0.9 were sequenced in the former study [9]. Since there is no chromosomal-scale of genome assembly for *M. coruscus*, the number of mis-assemblies were not documented for this species. For Wtdbg2 assemblies, the highest quality also resulted from the combination of NECAT-corrected and trimmed ONT reads, with the NG50 of 1.12Mb, like in the case of the Scaly-foot Snail above. However, the running time for *M. coruscus* was significantly inflated due to extensive computing required for the read error correction and trimming. Meanwhile, for Flye assemblies, the genome assembly from raw ONT reads was better than the assembly resulting from NECA-corrected and trimmed reads. However, neither of these two Flye assemblies had N50 values over 500Kb. This is rather different from the results in the Scaly-foot Snail assembly, and it may indicate the *M. coruscus* genome is too heterozygous or repetitive for Flye to be an effective assembler.

Among all of the *M. coruscus* assemblies generated in our benchmarking, the NextDenovo version exhibited the highest NG50 (3.40Mb) and BUSCO scores (Table 3, Supplementary Table S4). The MaSuRCA version resulted in the same ‘Complete’ BUSCO score, but with higher ‘Complete and Duplicated’ score (2.2% vs 1.7%) (Supplementary Table S4), suggesting that NextDenovo performs better in merging allelic contigs. Shasta was again the speediest, followed by Wtdbg2 and then Raven. Since the NextDenovo assembly was by far the most contiguous genome, this was selected for the downstream analysis. After three rounds of polishing with ONT reads, purging redundant haplotigs with purge_dups, and two rounds of error correction with Illumina reads using Pilon, the final N50 reached 2.54Mb, and the complete BUSCO score was 95.8% (duplicated BUSCO =1.7%). This is a dramatic improvement from the original published assembly (N50 = 898.3Kb and complete BUSCO score = 91.7%, and duplicated BUSCOs = 2.5%) [9]. A total of 72,541 gene models were annotated from this genome assembly, with the BUSCO score of the gene models being 92.3%. The number of gene models is rather high among published Mollusca genomes. A recent genome assembly of its congener *Mytilus galloprovincialis* annotated 60,338 genes, and the authors suggested there is significant variation in gene presence / absence among *Mytilus* species [39]. These results collectively indicate that *Mytilus* is gene-rich, and a similar high number of genes was also reported from another lamellibranch bivalve, the scallop *Pecten maximus* with 67,741 genes [40].

**Table 3.**
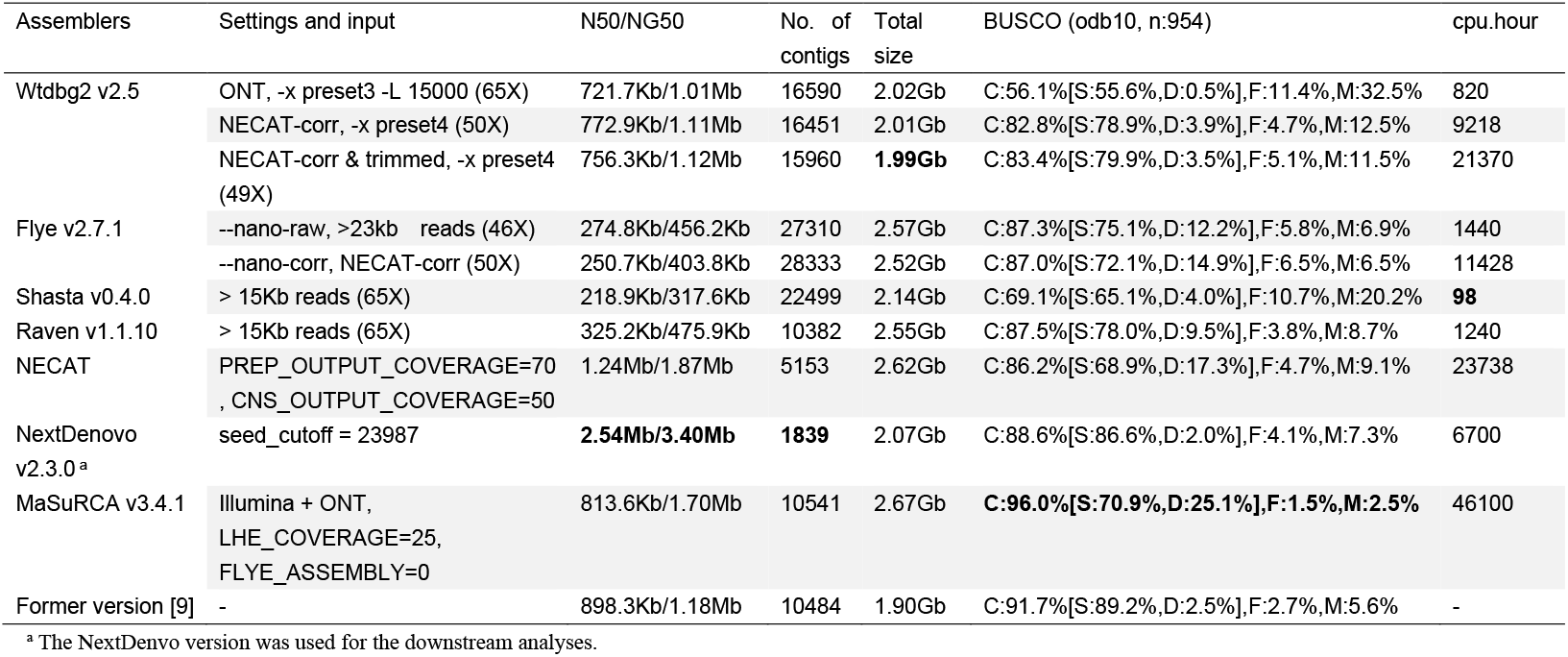
Assembly results for the *Mytilus coruscus* genome. The best performance was highlighted in bold. NG50 is the N50 value after normalization using the predicted genome size.

| Assemblers | Settings and input | N50/NG50 | No. of contigs | Total size | BUSCO (odb10, n:954) | cpu.hour |
| --- | --- | --- | --- | --- | --- | --- |
| Wtdbg2 v2.5 | ONT, -x preset3 -L 15000 (65X) | 721.7Kb/1.01Mb | 16590 | 2.02Gb | C:56.1%[S:55.6%,D:0.5%],F:11.4%,M:32.5% | 820 |
|  | NECAT-corr, -x preset4 (50X) | 772.9Kb/1.11Mb | 16451 | 2.01Gb | C:82.8%[S:78.9%,D:3.9%],F:4.7%,M:12.5% | 9218 |
|  | NECAT-corr & trimmed, -x preset4 (49X) | 756.3Kb/1.12Mb | 15960 | **1.99Gb** | C:83.4%[S:79.9%,D:3.5%],F:5.1%,M:11.5% | 21370 |
| Flye v2.7.1 | --nano-raw, >23kb reads (46X) | 274.8Kb/456.2Kb | 27310 | 2.57Gb | C:87.3%[S:75.1%,D:12.2%],F:5.8%,M:6.9% | 1440 |
|  | --nano-corr, NECAT-corr (50X) | 250.7Kb/403.8Kb | 28333 | 2.52Gb | C:87.0%[S:72.1%,D:14.9%],F:6.5%,M:6.5% | 11428 |
| Shasta v0.4.0 | > 15Kb reads (65X) | 218.9Kb/317.6Kb | 22499 | 2.14Gb | C:69.1%[S:65.1%,D:4.0%],F:10.7%,M:20.2% | **98** |
| Raven v1.1.10 | > 15Kb reads (65X) | 325.2Kb/475.9Kb | 10382 | 2.55Gb | C:87.5%[S:78.0%,D:9.5%],F:3.8%,M:8.7% | 1240 |
| NECAT | PREP_OUTPUT_COVERAGE=70, CNS_OUTPUT_COVERAGE=50 | 1.24Mb/1.87Mb | 5153 | 2.62Gb | C:86.2%[S:68.9%,D:17.3%],F:4.7%,M:9.1% | 23738 |
| NextDenovo v2.3.0<sup>a</sup> | seed_cutoff = 23987 | **2.54Mb/3.40Mb** | **1839** | 2.07Gb | C:88.6%[S:86.6%,D:2.0%],F:4.1%,M:7.3% | 6700 |
| MaSuRCA v3.4.1 | Illumina + ONT, LHE_COVERAGE=25, FLYE_ASSEMBLY=0 | 813.6Kb/1.70Mb | 10541 | 2.67Gb | **C:96.0%[S:70.9%,D:25.1%],F:1.5%,M:2.5%** | 46100 |
| Former version [9] | - | 898.3Kb/1.18Mb | 10484 | 1.90Gb | C:91.7%[S:89.2%,D:2.5%],F:2.7%,M:5.6% | - |

<sup>a</sup> The NextDenovo version was used for the downstream analyses.

### (c) Overall remark on the two genome assemblies

The genomes of *Chrysomallon squamiferum* and *Mytilus coruscus* represent two genomes with drastically different genomic features: the former genome is compact and the latter is relatively large; although both genomes are very heterozygous the latter is much more so. Genome assemblies with different assemblers resulted in various levels of trade-off between time, contiguity, and completeness. In general, Shasta, Wtdbg2, and Raven are very speedy and are the recommended assemblers when a quick check of the genomic features is desired instead of a high-quality assembly. Flye is not sensitive to the read accuracy, but Wtdbg2 always performed better with corrected and trimmed reads. NextDenovo was the highest performer in terms of genome contiguity (e.g., N50), but the genome completeness assessed by the BUSCO score was not the best, indicating the assembler likely failed to assemble some parts of the genome. We also found that QuickMerge can increase the genomic contiguity without introducing misassemblies, mismatches, or erroneous indels (Table 2). This is very impressive, since QuickMerge can be a cost-effective method for assembling a relatively high-quality genome. However, it should be noted that the BUSCO score after the QuickMerge is actually worsened, suggesting some parts of the genome have been lost during the consensus step.

**Table 2.**
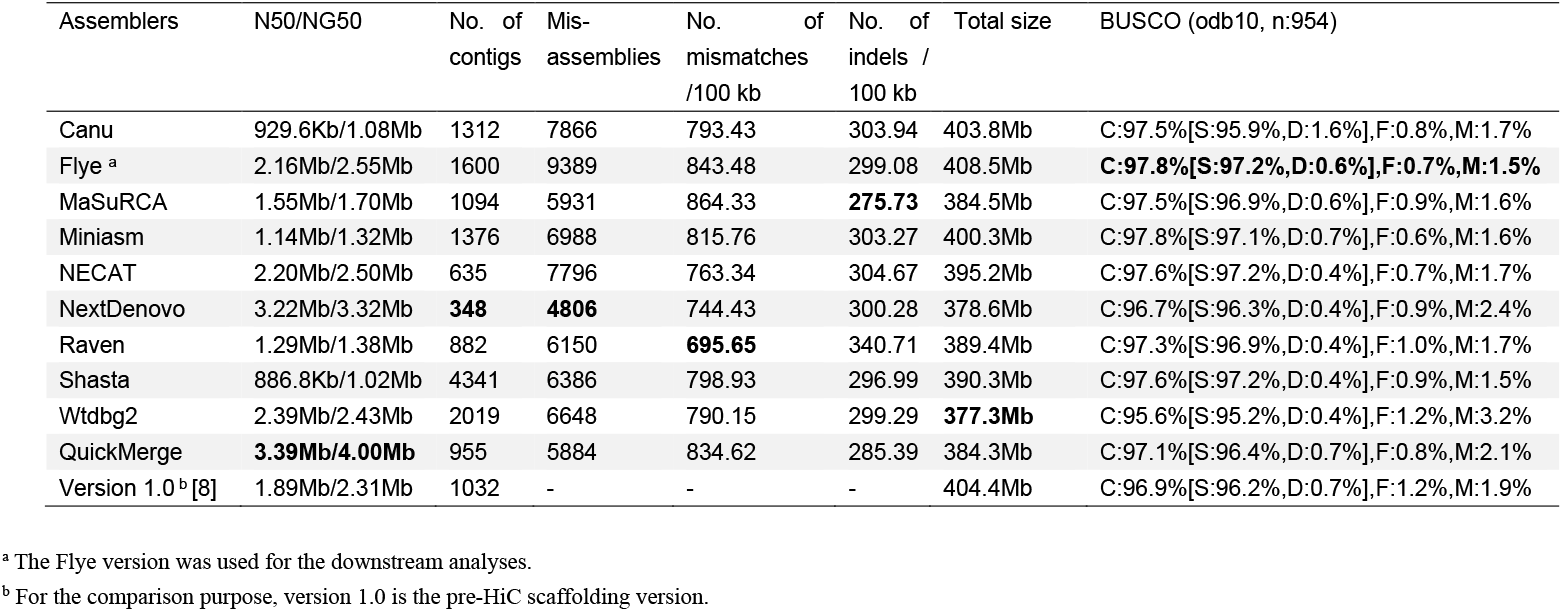
Assembly results for the Scaly-foot Snail *Chrysomallon squamiferum* genome from the nine assemblers and QuickMerge. The best performance was highlighted in bold. The input of the QuickMerge is the Flye assembly (the best BUSCO score) and MaSuRCA assembly (hybrid assembly). NG50 is the N50 value after normalization using the predicted genome size.

Regarding the input reads, our results demonstrate that using the longest 40-50x of ONT reads is recommended for assembling molluscan genomes. In the case of NextDenovo, the authors suggested the longest 45X of ONT reads as the optimised input; in the case of Flye, the authors suggested longest 40X of ONT reads. We found that including shorter reads in the assembly in most cases resulted in lower quality assemblies. Haplotig reduction via the purge_dups pipeline, which performs best in our experience (data not shown), or a similar approach is necessary for a genome assembly larger than the predicted genome size, because heterozygous genomic regions can inflate the assembly size. Also, the BUSCO score remains the same or even improved after the purge_dups pipeline (Supplementary Table S3), suggesting the heterozygous contigs have detrimental effects on the BUSCO score.

In general, of the 10 assemblers tested, Flye and Nextdenovo performed better than the rest overall. However, their performance was vastly different in the two species tested, with Flye performing the best in the *C. squamiferum* genome and NextDenovo in *M. coruscus*. With an extremely heterozygous genome like *M. coruscus*, NextDenovo likely performs better than Flye and is the recommended assembler with ONT reads. When the sequencing effort is limited, we also suggest using QuickMerge to merge at least two versions of the assembly in order to increase the genome continuousness without sacrificing too much assembly accuracy loss of genomic regions as reflected by a reduced BUSCO score.

### (d) Phylogenomic analyses on the available molluscan genomes

With these two updated genomes, we re-analysed the genome-level phylogeny of Mollusca including other publicly available high-quality molluscan genomes. Our pipeline recovered 5,388 orthologous genes and an alignment totalling 1,727,673 amino acid positions. All genes were sampled for at least 17/34 taxa with an average of 30 taxa sampled per alignment and 14.90% missing data overall in the resulting matrix. The resulting maximum likelihood tree exhibited maximum support at every node (Figure 1). Among the only three molluscan classes with high-quality genomes available at the time these analyses were performed, Cephalopoda was recovered sister to a clade comprising Bivalvia and Gastropoda, similar to previous studies [4, 8]. However, this result does not necessarily reflect sister relationships per se among clades, given the limited availability of taxa. Similarly, since only one decapodiform and one octopodiform was available, genomic insights of the relationships within Cephalopoda must await better taxon sampling in the future. In Bivalvia, the only significant difference with previously published phylogenies is the position of the order Arcida, which was previously recovered sister to the rest of Pteriomorpha [41] but here it was recovered as sister to Pectinida. Instead, our tree infers the split between Arcida and Pectinida within Pteriomorpha occurred later than the split between an Arcida/Pectinida clade with a Mytilida/Ostreida clade. The bivalve taxon sampling of high-quality genomes, however, continues to suffer from a heavy bias to the clade Pteriomorphia. Although in recent years a number of representatives of Imparidentia have been sequenced, all other major bivalve clades including Protobranchia, Paleoheterodonta, Archiheterodonta, and Anomalodesmata remain unrepresented. Since only two major clades have high-quality genomes, the relationships among major bivalve clades also remain a key topic of future genomic research. Within Gastropoda, the relationships among major subclass-level clades, as well as families, remained similar to previous phylogenomic trees [42]. A split between Patellogastropoda/Vetigastropoda/Neomphaliones and Caenogastropoda/Heterobranchia was seen, and, within the former clade, Neomphalida was sister to Vetigastropoda and this pair was in turn sister to Patellogastropoda. However, understanding of the internal relationships among gastropods continues to suffer from a lack of sufficient taxon sampling, such as the total lack of members of the subclass Neritimorpha. A mitochondrial genome-based phylogeny including all gastropod subclasses recovered Patellogastropoda sister to the rest of Gastropoda, which is in-line with evidence from fossils and morphology [43] but different from a phylogenomic study based on transcriptomes where it was recovered sister to Vetigastropoda [42]. Patellogastropods are also thought to suffer from long-branch attraction, and it is difficult to resolve this group’s phylogenetic position without a dense taxon sampling, as demonstrated by the mitogenome study where the position of Patellogastropoda was only reliably resolved when multiple genera were included in addition to *Lottia*. We hope more high-quality genome assemblies will be published at a faster pace in the near future, especially for currently under-sampled groups in Mollusca [44], using our benchmarking presented herein as a guide to achieving high efficiency.

**Figure 1.**
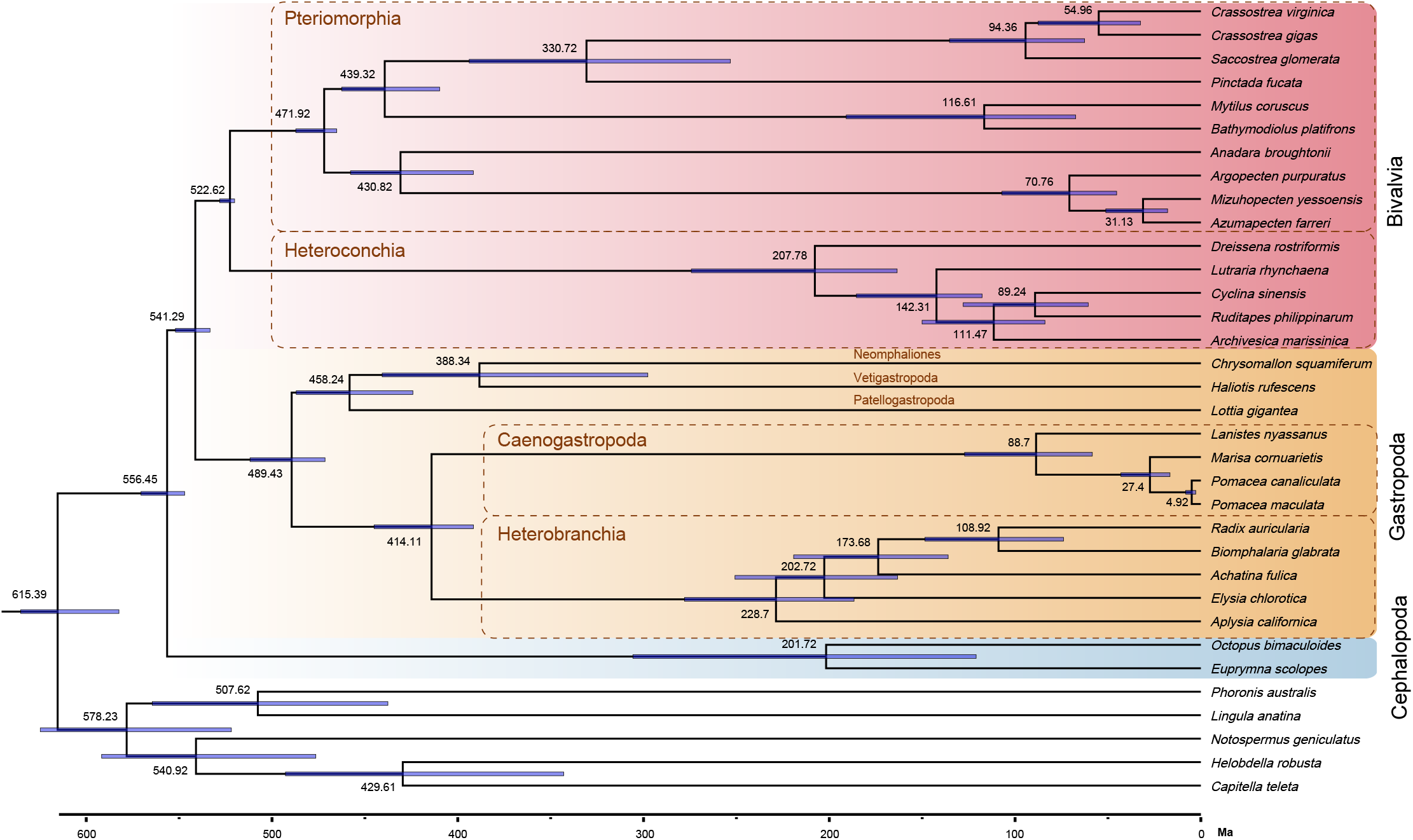
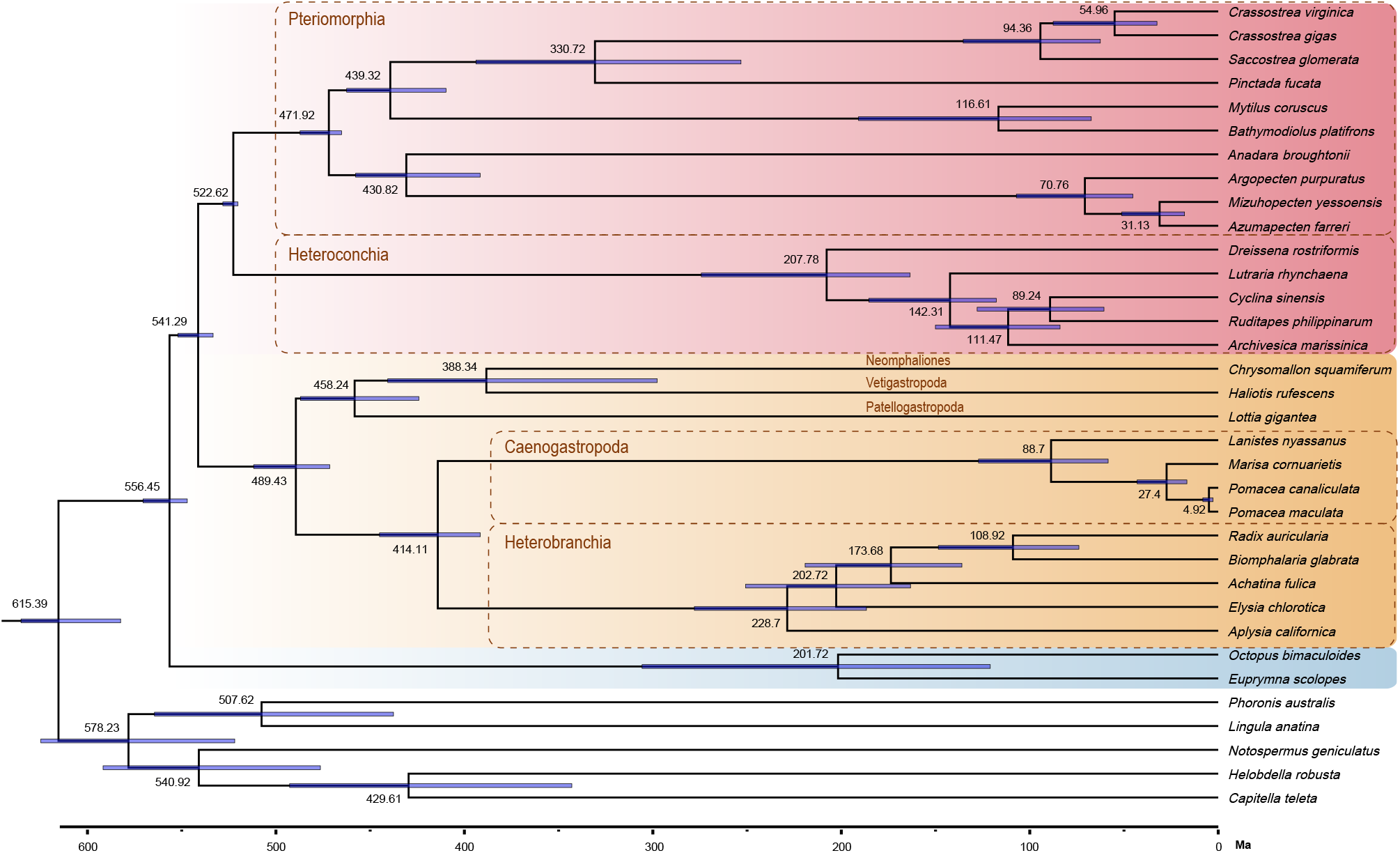
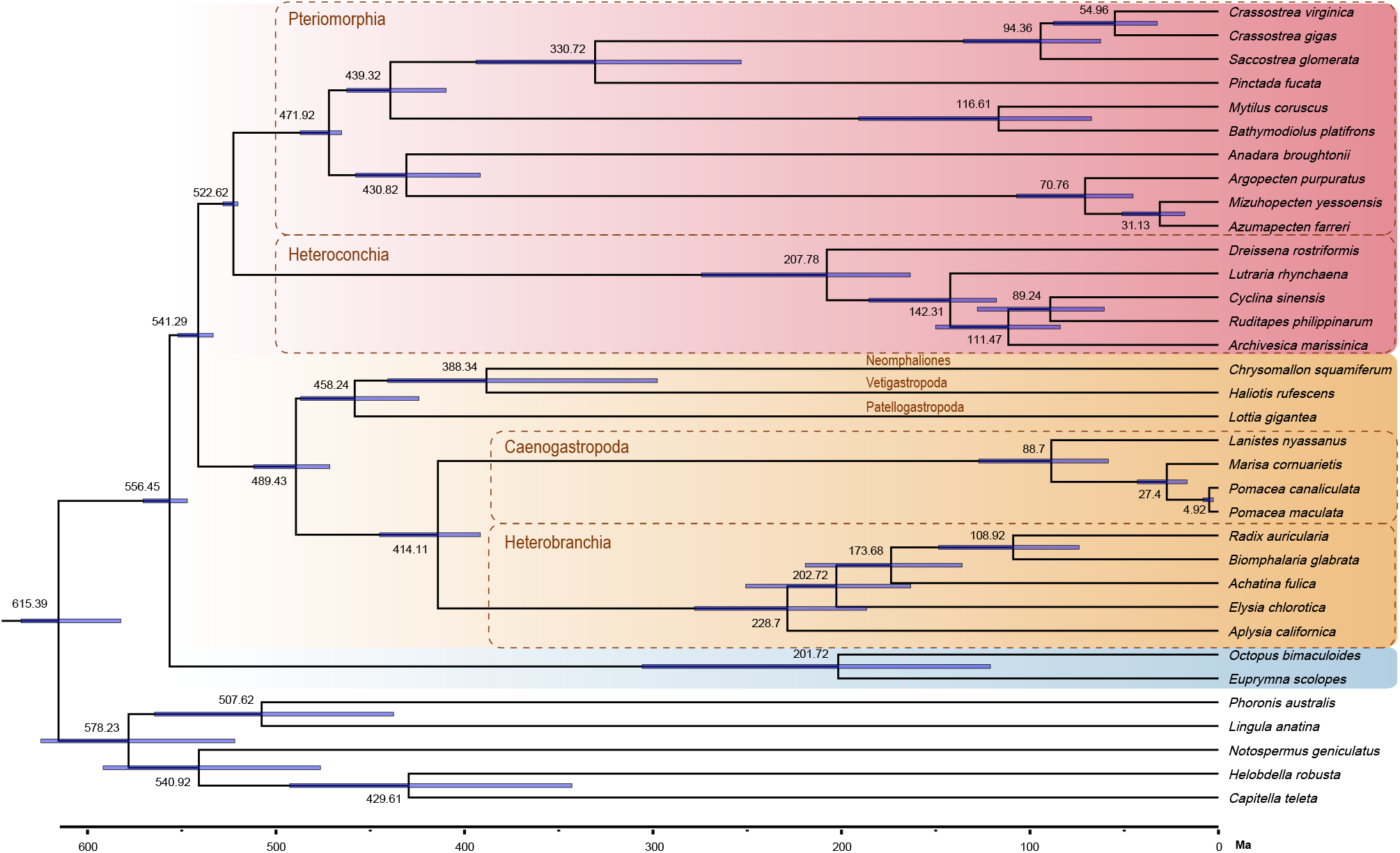
Evolutionary time tree of Mollusca and five other lophotrochozoans inferred from MCMCTree analysis. The error bar on each node indicates 95% confidence level. Background colours represent different molluscan classes: pink, Bivalvia; orange, Gastropoda; blue, Cephalopoda.

We found a positive, statistically significant, correlation (r = 0.342, *P* = 0.048) between BUSCO score and orthologue gene occupancy per genome used in the phylogenomic analysis (Figure 2). This indicates increasing the genome quality indeed leads to better coverage per orthologue group, thereby benefitting phylogenomic analyses in increasing the completeness of the data matrix that can be used. The two genomes newly updated herein, i.e. *C. squamiferum* and *M. coruscus*, have both higher BUSCO scores and orthologue gene occupancy compared to most of other published molluscan genomes, exemplifying that re-assembly of existing genomes using improved techniques is beneficial and useful. Compared with other molluscan genomes available in Gastropoda and Bivalvia, the two cephalopod genomes exhibited comparatively low BUSCO score and also orthologue gene occupancy. More optimised, higher-quality genome assemblies are required in Cephalopoda for a better coverage in the orthogroups, in order to improve the quality of phylogenomic analyses both within Cephalopoda and Mollusca.

**Figure 2.**
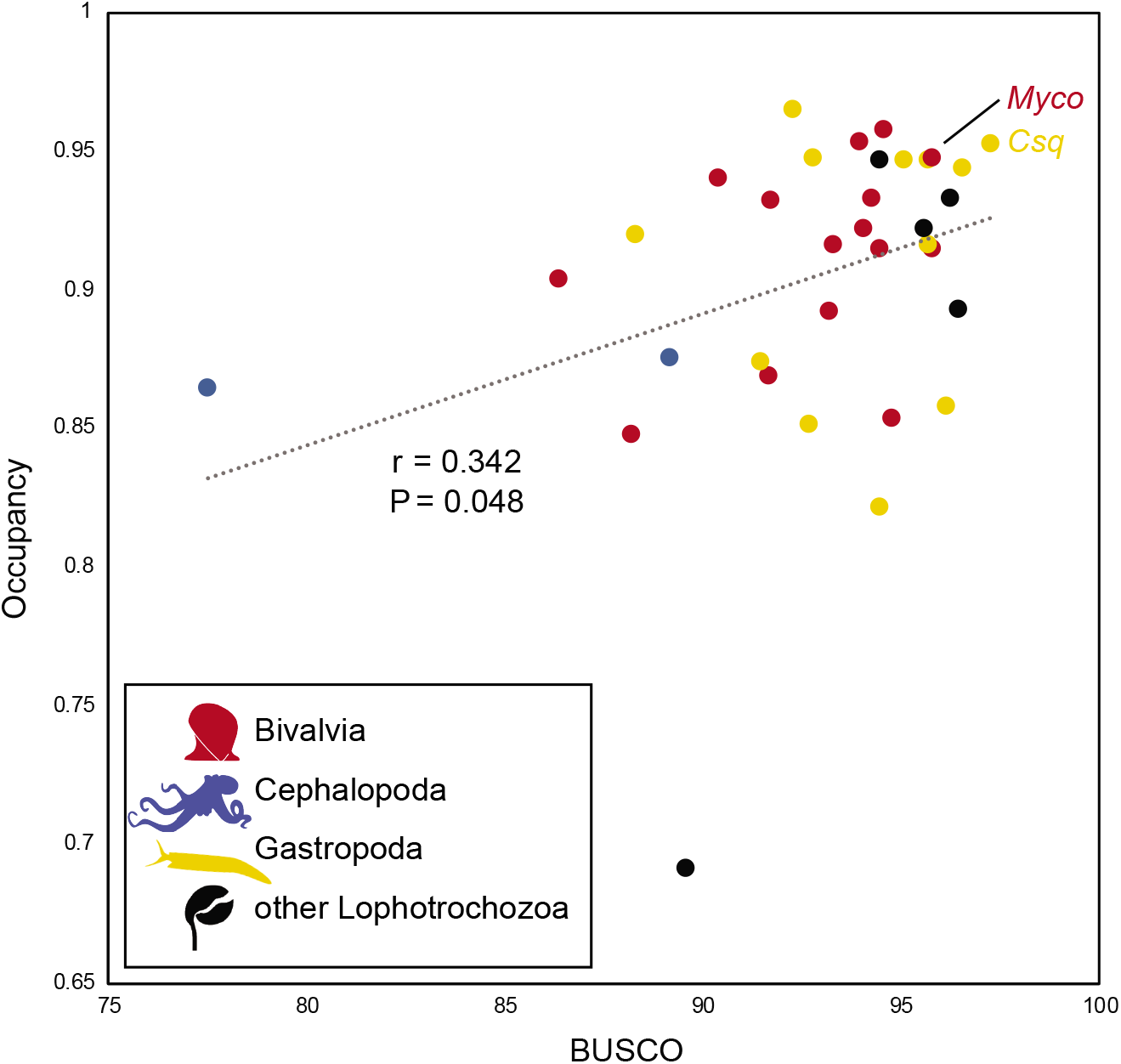
A correlation analysis between BUSCO score and orthologue gene occupancy per lophotrochozoan species with high-quality genomes available. Abbreviations: Csq, *Chrysomallon squamiferum* and Myco, *Mytilus coruscu*.

## Conclusions

We carried out benchmarking of various genome assemblers for the ONT using data from two molluscs, which suggested a 40-50X of ONT reads to be sufficient for achieving a high-quality genome assembly. Although different assemblers showed varying performances on different scores, overall Flye appears to be the best assembler for relatively simple genomes in Mollusca exemplified by *Chrysomallon squamiferum*, while NextDenovo performs the best for more complicated molluscan genomes exemplified by *Mytilus coruscus*. Increasing the genome assembly quality is beneficial to various downstream analyses, for instance, by increasing the completeness of the sampling matrices in the phylogenetic analysis. These results may also be applicable to other important yet neglected groups of marine invertebrates, such as polychaetes, brachiopods, nemerteans, and other lophotrochozoans, which may share similar genomic features with molluscs. In the future, it is necessary to assess the genome assembly quality with ultra-long ONT reads and also PacBio HiFi reads, which are newly available techniques with very little sequencing errors, and a comprehensive comparison between these two sequencing techniques are also needed.

## Acknowledgments

We thank Dr. Angus Davison and Dr. Maurine Neiman for organising the Theo Murphy international scientific meeting ‘Pearls of wisdom: synergising leadership and expertise in molluscan genomics’ and for curating and editing this special issue. This work was supported by the Young Taishan Scholars Program of Shandong Province.

## Supplementary material

Supplementary Information contains all commands used for genome assemblies used in the present study.

**Supplementary Table S1.** List of currently available lophotrochozoan genomes and their completeness. Genomes with a BUSCO score of the predicted gene models over 90% are highlighted in yellow, and genomes with a BUSCO score over 80% are highlighted in orange. Genomes published without annotated gene models were excluded.

**Supplementary Table S2.** Detailed assembly results for the Scaly-foot Snail (*Chrysomallon squamiferum*) genome, with the best performance from each assembler being highlighted in yellow. Corr, corrected reads, trim, corrected and trimmed reads. NG50 represents the N50 value after normalization using the predicted genome size.

**Supplementary Table S3.** Detailed Assembly results for the Scaly-foot Snail (*Chrysomallon squamiferum*) genome from the nine assemblers and QuickMerger. The best performance being highlighted in yellow. The input of QuickMerger is the Flye assembly (the best BUSCO score) and MaSuRCA assembly (hybrid assembly). NG50 represents the N50 value after normalization using the predicted genome size.

**Supplementary Table S4.** Detailed assembly results for the Hard-shelled Mussel (*Mytilus coruscus*) genome, with the best performance from each assembler being highlighted in yellow. Corr, corrected reads, trim, corrected and trimmed reads. NG50 represents the N50 value after normalization using the predicted genome size.

## Ethics

No human or animals were used in this study. All of the raw sequencing data were generated from the former studies.

## Data Accessibility

The raw ONT reads basecalled in this study were deposit in the NCBI SRA database with the accession number SRR12763791. The assembled genomes, files related to the OrthoFinder and phylogenomic analyses, the gene model annotations of both species and the InterProScan output, were deposited to Dryad with the DOI: 10.5061/dryad.w6m905qns. Commands used for genome assemblies are provided in the Supplementary Information.

## Authors’ Contributions

J.S, K.M.K and C.C. conceived and designed this research. J.S. and R. L. assembled the genomes and compared different assemblies. K.M.K. performed the OrthoFinder and phylogenomic analyses, and annotated the gene models. J.S. and R.L. performed the gene family analysis. J.S. C.C., K.M.K. and J.D.S. prepared and revised the manuscript.

## References

1 Matz, M. V. 2018 Fantastic beasts and how to sequence them: ecological genomics for obscure model organisms. Trends in Genetics. 34, 121–132. (10.1016/j.tig.2017.11.002)

2 Takeuchi, T. 2017 Molluscan genomics: implications for biology and aquaculture. Current Molecular Biology Reports. (10.1007/s40610-017-0077-3)

3 Sokolow, S. H., Huttinger, E., Jouanard, N., Hsieh, M. H., Lafferty, K. D., Kuris, A. M., Riveau, G., Senghor, S., Thiam, C., N’Diaye, A., et al. 2015 Reduced transmission of human schistosomiasis after restoration of a native river prawn that preys on the snail intermediate host. Proc. Natl. Acad. Sci. 112, 9650–9655. (10.1073/pnas.1502651112)

4 Kocot, K. M., Cannon, J. T., Todt, C., Citarella, M. R., Kohn, A. B., Meyer, A., Santos, S. R., Schander, C., Moroz, L. L., Lieb, B., et al. 2011 Phylogenomics reveals deep molluscan relationships. Nature. 477, 452–456. (10.1038/nature10382)

5 Kocot, K. M., Poustka, A. J., Stöger, I., Halanych, K. M., Schrödl, M. 2020 New data from Monoplacophora and a carefully-curated dataset resolve molluscan relationships. Sci. Rep. 10, 101. (10.1038/s41598-019-56728-w)

6 Chen, Y., Nie, F., Xie, S.-Q., Zheng, Y.-F., Bray, T., Dai, Q., Wang, Y.-X., Xing, J.-f., Huang, Z.-J., Wang, D.-P., et al. 2020 Fast and accurate assembly of Nanopore reads via progressive error correction and adaptive read selection. bioRxiv. 2020.2002.2001.930107. (10.1101/2020.02.01.930107)

7 Shafin, K., Pesout, T., Lorig-Roach, R., Haukness, M., Olsen, H. E., Bosworth, C., Armstrong, J., Tigyi, K., Maurer, N., Koren, S., et al. 2020 Nanopore sequencing and the Shasta toolkit enable efficient de novo assembly of eleven human genomes. Nature Biotechnology. 38, 1044–1053. (10.1038/s41587-020-0503-6)

8 Sun, J., Chen, C., Miyamoto, N., Li, R., Sigwart, J. D., Xu, T., Sun, Y., Wong, W. C., Ip, J. C. H., Zhang, W., et al. 2020 The Scaly-foot Snail genome and implications for the origins of biomineralised armour. Nature Communications. 11, 1657. (10.1038/s41467-020-15522-3)

9 Li, R., Zhang, W., Lu, J., Zhang, Z., Mu, C., Song, W., Migaud, H., Wang, C., Bekaert, M. 2020 The whole-genome sequencing and hybrid assembly of *Mytilus coruscus*. Frontiers in Genetics. 11, (10.3389/fgene.2020.00440)

10 Wood, D. E., Lu, J., Langmead, B. 2019 Improved metagenomic analysis with Kraken 2. Genome Biology. 20, 257. (10.1186/s13059-019-1891-0)

11 Ranallo-Benavidez, T. R., Jaron, K. S., Schatz, M. C. 2020 GenomeScope 2.0 and Smudgeplot for reference-free profiling of polyploid genomes. Nature Communications. 11, 1432. (10.1038/s41467-020-14998-3)

12 Koren, S., Walenz, B. P., Berlin, K., Miller, J. R., Bergman, N. H., Phillippy, A. M. 2017 Canu: scalable and accurate long-read assembly via adaptive k-mer weighting and repeat separation. Genome Research. 27, 722–736. (10.1101/gr.215087.116)

13 Kolmogorov, M., Yuan, J., Lin, Y., Pevzner, P. A. 2019 Assembly of long, error-prone reads using repeat graphs. Nature Biotechnology. 37, 540–546. (10.1038/s41587-019-0072-8)

14 Ruan, J., Li, H. 2019 Fast and accurate long-read assembly with wtdbg2. Nature Methods. 17, 155–158. (10.1038/s41592-019-0669-3)

15 Li, H. 2016 Minimap and miniasm: fast mapping and de novo assembly for noisy long sequences. Bioinformatics. 32, 2103–2110. (10.1093/bioinformatics/btw152)

16 Vaser, R., Šikić, M. 2020 Raven: a de novo genome assembler for long reads. bioRxiv. 2020.2008.2007.242461. (10.1101/2020.08.07.242461)

17 Zimin, A. V., Puiu, D., Luo, M.-C., Zhu, T., Koren, S., Marçais, G., Yorke, J. A., Dvořák, J., Salzberg, S. L. 2017 Hybrid assembly of the large and highly repetitive genome of Aegilops tauschii, a progenitor of bread wheat, with the MaSuRCA mega-reads algorithm. Genome Research. 27, 787–792. (10.1101/gr.213405.116)

18 Chakraborty, M., Baldwin-Brown, J. G., Long, A. D., Emerson, J. J. 2016 Contiguous and accurate de novo assembly of metazoan genomes with modest long read coverage. Nucleic Acids Research. 44, e147–e147. (10.1093/nar/gkw654)

19 Schmidt, M. H., Vogel, A., Denton, A. K., Istace, B., Wormit, A., van de Geest, H., Bolger, M. E., Alseekh, S., Maß, J., Pfaff, C., et al. 2017 De novo assembly of a new *Solanum pennellii* accession using Nanopore sequencing. The Plant Cell. 29, 2336–2348. (10.1105/tpc.17.00521)

20 Guan, D., McCarthy, S. A., Wood, J., Howe, K., Wang, Y., Durbin, R. 2020 Identifying and removing haplotypic duplication in primary genome assemblies. Bioinformatics. 36, 2896–2898. (10.1093/bioinformatics/btaa025)

21 Walker, B. J., Abeel, T., Shea, T., Priest, M., Abouelliel, A., Sakthikumar, S., Cuomo, C. A., Zeng, Q., Wortman, J., Young, S. K., et al. 2014 Pilon: An integrated tool for comprehensive microbial variant detection and genome assembly improvement. PLoS ONE. 9, e112963. (10.1371/journal.pone.0112963)

22 Seppey, M., Manni, M., Zdobnov, E. M. 2019 BUSCO: Assessing genome assembly and annotation completeness. In Gene Prediction: Methods and Protocols. (ed.^eds. M. Kollmar), pp. 227–245. New York, NY: Springer New York.

23 Gurevich, A., Saveliev, V., Vyahhi, N., Tesler, G. 2013 QUAST: quality assessment tool for genome assemblies. Bioinformatics. 29, 1072–1075. (10.1093/bioinformatics/btt086)

24 Emms, D. M., Kelly, S. 2019 OrthoFinder: phylogenetic orthology inference for comparative genomics. Genome Biology. 20, 238. (10.1186/s13059-019-1832-y)

25 Katoh, K., Misawa, K., Kuma, K. i., Miyata, T. 2002 MAFFT: a novel method for rapid multiple sequence alignment based on fast Fourier transform. Nucleic Acids Research. 30, 3059–3066. (10.1093/nar/gkf436)

26 Di Franco, A., Poujol, R., Baurain, D., Philippe, H. 2019 Evaluating the usefulness of alignment filtering methods to reduce the impact of errors on evolutionary inferences. BMC Evolutionary Biology. 19, 21. (10.1186/s12862-019-1350-2)

27 Criscuolo, A., Gribaldo, S. 2010 BMGE (Block Mapping and Gathering with Entropy): a new software for selection of phylogenetic informative regions from multiple sequence alignments. BMC Evolutionary Biology. 10, 210. (10.1186/1471-2148-10-210)

28 Price, M. N., Dehal, P. S., Arkin, A. P. 2010 FastTree 2 - Approximately maximum-likelihood trees for large alignments. PLoS ONE. 5, e9490. (10.1371/journal.pone.0009490)

29 Minh, B. Q., Schmidt, H. A., Chernomor, O., Schrempf, D., Woodhams, M. D., von Haeseler, A., Lanfear, R. 2020 IQ-TREE 2: New models and efficient methods for phylogenetic inference in the genomic era. Mol. Biol. Evol. 37, 1530–1534. (10.1093/molbev/msaa015)

30 dos Reis, M., Thawornwattana, Y., Angelis, K., Telford, Maximilian J., Donoghue, Philip C. J., Yang, Z. 2015 Uncertainty in the timing of origin of animals and the limits of precision in molecular timescales. Current Biology. 25, 2939–2950. (10.1016/j.cub.2015.09.066)

31 Bieler, R., Mikkelsen, P. M., Collins, T. M., Glover, E. A., González, V. L., Graf, D. L., Harper, E. M., Healy, J., Kawauchi, G. Y., Sharma, P. P., et al. 2014 Investigating the Bivalve Tree of Life – an exemplar-based approach combining molecular and novel morphological characters. Invertebrate Systematics. 28, 32–115, 184.

32 Stoger, I., Sigwart, J. D., Kano, Y., Knebelsberger, T., Marshall, B. A., Schwabe, E., Schrödl, M. 2013 The continuing debate on deep molluscan phylogeny: Evidence for Serialia (Mollusca, Monoplacophora + Polyplacophora). BioMed Research International. 2013, 18. (10.1155/2013/407072)

33 Sun, J., Mu, H., Ip, J. C. H., Li, R., Xu, T., Accorsi, A., Sánchez Alvarado, A., Ross, E., Lan, Y., Sun, Y., et al. 2019 Signatures of divergence, invasiveness, and terrestrialization revealed by four apple snail genomes. Mol. Biol. Evol. 36, 1507–1520. (10.1093/molbev/msz084)

34 Tillier, S. 1996 Phylogenetic relationships of the pulmonate gastropods from rRNA sequences, and tempo and age of the stylommatophoran radiation. Origin and evolutionary radiation of the Mollusca. 267–284.

35 Benton, M., Donoghue, P., Asher, R. 2009 Calibrating and constraining molecular clocks. The timetree of life. 35–86.

36 Jörger, K. M., Stöger, I., Kano, Y., Fukuda, H., Knebelsberger, T., Schrödl, M. 2010 On the origin of Acochlidia and other enigmatic euthyneuran gastropods, with implications for the systematics of Heterobranchia. BMC Evolutionary Biology. 10, 323. (10.1186/1471-2148-10-323)

37 Benton, M. J., Donoghue, P. C., Asher, R. J., Friedman, M., Near, T. J., Vinther, J. 2015 Constraints on the timescale of animal evolutionary history. Palaeontologia Electronica. 18, 1–106.

38 Tyson, J. R., O’Neil, N. J., Jain, M., Olsen, H. E., Hieter, P., Snutch, T. P 2018 MinION-based long-read sequencing and assembly extends the *Caenorhabditis elegans* reference genome. Genome Research. 28, 266–274. (10.1101/gr.221184.117)

39 Gerdol, M., Moreira, R., Cruz, F., Gómez-Garrido, J., Vlasova, A., Rosani, U., Venier, P., Naranjo-Ortiz, M. A., Murgarella, M., Balseiro, P., et al. 2019 Massive gene presence/absence variation in the mussel genome as an adaptive strategy: first evidence of a pan-genome in Metazoa. bioRxiv. 781377. (10.1101/781377)

40 Kenny, N. J., McCarthy, S. A., Dudchenko, O., James, K., Betteridge, E., Corton, C., Dolucan, J., Mead, D., Oliver, K., Omer, A. D., et al. 2020 The gene-rich genome of the scallop Pecten maximus. GigaScience. 9, (10.1093/gigascience/giaa037)

41 Lemer, S., González, V. L., Bieler, R., Giribet, G. 2016 Cementing mussels to oysters in the pteriomorphian tree: a phylogenomic approach. Proc. Royal Soc. B. 283, 20160857. (doi:10.1098/rspb.2016.0857)

42 Cunha, T. J., Giribet, G. 2019 A congruent topology for deep gastropod relationships. Proc. Royal Soc. B. 286, 20182776. (doi:10.1098/rspb.2018.2776)

43 Uribe, J. E., Irisarri, I., Templado, J., Zardoya, R. 2019 New patellogastropod mitogenomes help counteracting long-branch attraction in the deep phylogeny of gastropod mollusks. Molecular Phylogenetics and Evolution. 133, 12–23. (10.1016/j.ympev.2018.12.019)

44 Horn, K. M., Anderson, F. E. 2020 Spiralian Genomes Reveal Gene Family Expansions Associated with Adaptation to Freshwater. Journal of Molecular Evolution. (10.1007/s00239-020-09949-x)

