## Supplementary Information for "Benchmarking Oxford Nanopore read assemblers for high-quality molluscan genomes"

***The commands used in this study.***

\## Remove Bacteria contaminated Illumina reads by Kraken2

/home/share/kraken2-2.0.8-beta/kraken2 -db /home/share/kraken2-2.0.8-beta/minikraken2_v1_8GB --threads 40 --paired SFG1.fq SFG2.fq --classified-out SFG_Bac#.fq --unclassified-out SFG_Host#.fq

#Remove the symbiont contaminated ONT reads

minimap2 -ax map-ont -t 40 ~/Data/SFG/Symbiont_genome/Bac.fasta SFG_HAC_3Kb.fq | samtools fastq -n -f 4 - > SFG_HAC_3Kb_BacFree.fq

#wtdbg2 assembly

~/App/wtdbg2/wtdbg2 -i ../Flye_5Kb_canu_trimm_ass/SFG_HA.trimmedReads.fasta.gz -t 40 -o SFG_HAC_canuTrim

~/App/wtdbg2/wtpoa-cns -t 40 -i SFG_HAC_canuTrim.ctg.lay.gz -fo SFG_HAC_canuTrim.ctg.lay.fa

\# Flye assembly and polishing

flye --nano-raw ../SFG_HAC_3Kb_BacFree.fq -g 0.3566g -o flye_ONT_HAC_3kb -t 40

flye --polish-target ../SFG_10Kb_HAC.fa --nanopore-raw ../SFG_HAC_3Kb_BacFree.fq --iterations 3 --out-dir flye_po_R3 --threads 40

\# minimap + miniasm assembly

minimap2 -X -t 20 -x ava-ont ../SFGB1_clean_3Kb.fq ../SFGB1_clean_3Kb.fq > reads.paf

miniasm -f ../SFGB1_clean_3Kb.fq reads.paf > SFG_default.gfa

awk '/^S/{print ">"$2"\n"$3}' SFG_default.gfa | fold > SFG_default.fa

\# canu assembly

~/App/canu-2.0/Linux-amd64/bin/canu -fast -p SFG_HA -d SFG_HA genomeSize=0.3566g -nanopore-raw SFG_HAC_3Kb_BacFree.fq corOutCoverage=200 corMhapSensitivity=normal correctedErrorRate=0.105 minReadLength=5000 useGrid=true gridOptions=--partition=oces

#NECAT assembly

necat.pl config SFG_HAC_config.txt

necat.pl correct SFG_HAC_config.txt

necat.pl assemble SFG_HAC_config.txt

necat.pl bridge SFG_HAC_config.txt

\# the configure file

PROJECT=SFG_HAC

ONT_READ_LIST=readlist.txt

GENOME_SIZE=356600000

THREADS=40

MIN_READ_LENGTH=5000

PREP_OUTPUT_COVERAGE=70

OVLP_FAST_OPTIONS=-n 500 -z 20 -b 2000 -e 0.5 -j 0 -u 1 -a 1000

OVLP_SENSITIVE_OPTIONS=-n 500 -z 10 -e 0.5 -j 0 -u 1 -a 1000

CNS_FAST_OPTIONS=-a 2000 -x 4 -y 12 -l 1000 -e 0.5 -p 0.8 -u 0

CNS_SENSITIVE_OPTIONS=-a 2000 -x 4 -y 12 -l 1000 -e 0.5 -p 0.8 -u 0

TRIM_OVLP_OPTIONS=-n 100 -z 10 -b 2000 -e 0.5 -j 1 -u 1 -a 400

ASM_OVLP_OPTIONS=-n 100 -z 10 -b 2000 -e 0.5 -j 1 -u 0 -a 400

NUM_ITER=2

CNS_OUTPUT_COVERAGE=50

CLEANUP=1

USE_GRID=false

GRID_NODE=0

GRID_OPTIONS=

SMALL_MEMORY=0

FSA_OL_FILTER_OPTIONS=

FSA_ASSEMBLE_OPTIONS=

FSA_CTG_BRIDGE_OPTIONS=

POLISH_CONTIGS=true

#NextDenovo configure file

[General]

job_type = local

job_prefix = nextDenovo

task = all # 'all', 'correct', 'assemble'

rewrite = yes # yes/no

deltmp = yes

rerun = 3

parallel_jobs = 4

input_type = raw

input_fofn = ./input.fofn

workdir = ./01_rundir

[correct_option]

read_cutoff = 1k

seed_cutoff = 11653

blocksize = 1g

pa_correction = 2

seed_cutfiles = 2

sort_options = -m 1g -t 2 -k 50

minimap2_options_raw = -x ava-ont -t 10

correction_options = -p 15

[assemble_option]

minimap2_options_cns = -x ava-ont -t 10 -k17 -w17

nextgraph_options = -a 1

#Raven assembly

~/App/raven/build/bin/raven ../SFG_HAC_10Kb_BacFree.fa.gz -t 40 -p 1 >Raven_10Kb.fasta 2> log.txt

#Shasta assembly

sudo ~/App/shata_v0.4.0/shasta-Linux-0.4.0 --input ../Mcor_ONT_10kb.fa --memoryMode filesystem --memoryBacking 2M

#QuickMerge

delta-filter -r -q -l 10000 SFG_hybrid.delta > SFG_hybrid.rq.delta

quickmerge -d SFG_hybrid.rq.delta -q MaSu_HAC35x_FlyeP3_PD_Pilon2.fasta -r SFG_Flye_HAC10kb_P2_PD_pilon2.fasta -hco 5.0 -c 1.5 -l 2164900 -ml 10000 -p SFG_QM

\# Purge_dup version 1.2.3

minimap2 -x map-ont SFG_FlyepolishedR3.fasta SFG_ONT_3Kb_BacFree.fq -t 40 > reads.paf

pbcstat *.paf

calcuts PB.stat -l13 -m51 -u144 > cutoffs 2>calcults.log

split_fa SFG_FlyepolishedR3.fasta > polished_3.fasta.split

minimap2 -x asm5 -DP polished_3.fasta.split polished_3.fasta.split -t 40 >split.paf

purge_dups -2 -T cutoffs -c PB.base.cov split.paf >dups.bed 2>purge_dups.log

/get_seqs dups.bed SFG_FlyepolishedR3.fasta

#or with Illumina reads

bowtie2-build -f SFG_canu_flyeP3.fasta SFG_canu --threads 40

bowtie2 -p 40 --maxins 800 -x SFG_canu -1 SFG_trim_1.fq -2 SFG_trim_2.fq 1>PE.sam 2>SE.err

samtools view -bS PE.sam >PE.bam -@ 20

~/App/purge_dups/bin/ngscstat PE.bam

~/App/purge_dups/bin/calcuts TX.stat -l13 -m51 -u144 > cutoffs 2>calcults.log

~/App/purge_dups/bin/split_fa SFG_canu_flyeP3.fasta > polished_3.fasta.split

minimap2 -x asm5 -DP polished_3.fasta.split polished_3.fasta.split -t 40 > polished_3.fasta.split.paf

~/App/purge_dups/bin/purge_dups -2 -T cutoffs -c TX.base.cov polished_3.fasta.split.paf > dups.bed 2> purge_dups.log

#Pilon error correction

bowtie2-build -f canu_FlyeP3_PD2_pilon1.fa SFS --threads 40

bowtie2 -p 40 -D 20 -R 2 -N 1 -L 18 -i S,1,0.50 --maxins 1200 -x SFS -1 SFG_clean_1.fq -2

SFG_clean_2.fq 1>SFSPE500.sam 2> SFSPE500.err

grep -E "@|NM:" SFSPE500.sam | grep -v "XS:" > SFSPE500_uniq.sam

samtools view -bS SFSPE500_uniq.sam > SFSPE500_uniq.bam -@ 40

samtools sort SFSPE500_uniq.bam -m 5G -@ 10 -o SFSPE500_uniq_sorted.bam

java -jar ~/App/picard/picard.jar MarkDuplicates I= SFSPE500_uniq_sorted.bam O= SFSPE500_uniq_sorted_dedupe.bam METRICS_FILE=metrics.txt

samtools index SFSPE500_uniq_sorted_dedupe.bam

java -Xmx180G -jar ~/App/pilon-1.23/pilon-1.23.jar --genome canu_FlyeP3_PD2_pilon1.fa --frags SFSPE500_uniq_sorted_dedupe.bam --diploid --threads 40

#QUAST analysis

quast.py ./canu/Canu.fasta ./flye/Flye.fasta ./masurca/MaSuRCA.fasta ./miniasm/Miniasm.fasta ./necat/NECAT.fasta ./nextdenovo/NextDenovo.fasta

./raven/Raven.fasta ./shasta/Shasta.fasta ./wtdbg2/Wtdbg2.fasta ./quickmerger/QuickMerger.fasta -r ./Csqv1.1/Csq_v1.1.fa -t 40

#repeatmodeler and repeatmasker

BuildDatabase -name Mcor_v2.0 ../Mcor_v2.0.fasta

~/App/RepeatModeler-2.0.1/RepeatModeler -database Mcor_v2.0 -pa 40

RepeatMasker -species all -pa 8 -div 30 Mcor_v2.0.fasta

RepeatMasker -lib /home/sunj/Data/Mcoru/Annotations/RepeatModeler/Mcor_v2.0-families.fa -pa 10 -div 30 Mcor_v2.0.fasta.masked

\# Braker for Augustus training

~/App/BRAKER-2.1.5/scripts/braker.pl --genome=SFG_Flye_HAC10kb_P2_PD_pilon2.fasta.masked.masked --species=Chrysomallon --bam=all.sorted.bam --cores 40

\# maker configure file

#-----Genome (these are always required)

genome=SFG_Flye_HAC10kb_P2_PD_pilon2.fasta #genome sequence (fasta file or fasta embeded in GFF3 file)

organism_type=eukaryotic #eukaryotic or prokaryotic. Default is eukaryotic

#-----EST Evidence (for best results provide a file for at least one)

est=Trinity_all_0.97.fasta #set of ESTs or assembled mRNA-seq in fasta format

altest= #EST/cDNA sequence file in fasta format from an alternate organism

est_gff= #aligned ESTs or mRNA-seq from an external GFF3 file

altest_gff= #aligned ESTs from a closly relate species in GFF3 format

#-----Protein Homology Evidence (for best results provide a file for at least one)

protein=Mollu_Prot_50AA_0.95.fa #protein sequence file in fasta format (i.e. from mutiple organisms)

protein_gff= #aligned protein homology evidence from an external GFF3 file

#-----Repeat Masking (leave values blank to skip repeat masking)

model_org=all #select a model organism for RepBase masking in RepeatMasker

rmlib=./SFG_flye-families.fa #provide an organism specific repeat library in fasta format for RepeatMasker

repeat_protein=~/App/maker-3.01.03/data/te_proteins.fasta #provide a fasta file of transposable element proteins for RepeatRunner

rm_gff= #pre-identified repeat elements from an external GFF3 file

prok_rm=0 #forces MAKER to repeatmask prokaryotes (no reason to change this), 1 = yes, 0 = no

softmask=1 #use soft-masking rather than hard-masking in BLAST (i.e. seg and dust filtering)

#-----Gene Prediction

snaphmm= #SNAP HMM file

gmhmm= #GeneMark HMM file

augustus_species=Chrysomallon #Augustus gene prediction species model

fgenesh_par_file= #FGENESH parameter file

pred_gff= #ab-initio predictions from an external GFF3 file

model_gff= #annotated gene models from an external GFF3 file (annotation pass-through)

run_evm=1 #run EvidenceModeler, 1 = yes, 0 = no

est2genome=1 #infer gene predictions directly from ESTs, 1 = yes, 0 = no

protein2genome=1 #infer predictions from protein homology, 1 = yes, 0 = no

trna=0 #find tRNAs with tRNAscan, 1 = yes, 0 = no

snoscan_rrna= #rRNA file to have Snoscan find snoRNAs

snoscan_meth= #-O-methylation site fileto have Snoscan find snoRNAs

unmask=0 #also run ab-initio prediction programs on unmasked sequence, 1 = yes, 0 = no

allow_overlap= #allowed gene overlap fraction (value from 0 to 1, blank for default)

#-----Other Annotation Feature Types (features MAKER doesn't recognize)

other_gff= #extra features to pass-through to final MAKER generated GFF3 file

#-----External Application Behavior Options

alt_peptide=C #amino acid used to replace non-standard amino acids in BLAST databases

cpus=20 #max number of cpus to use in BLAST and RepeatMasker (not for MPI, leave 1 when using MPI)

#-----MAKER Behavior Options

max_dna_len=100000 #length for dividing up contigs into chunks (increases/decreases memory usage)

min_contig=10000 #skip genome contigs below this length (under 10kb are often useless)

pred_flank=200 #flank for extending evidence clusters sent to gene predictors

pred_stats=0 #report AED and QI statistics for all predictions as well as models

AED_threshold=1 #Maximum Annotation Edit Distance allowed (bound by 0 and 1)

min_protein=50 #require at least this many amino acids in predicted proteins

alt_splice=0 #Take extra steps to try and find alternative splicing, 1 = yes, 0 = no

always_complete=1 #extra steps to force start and stop codons, 1 = yes, 0 = no

map_forward=0 #map names and attributes forward from old GFF3 genes, 1 = yes, 0 = no

keep_preds=0 #Concordance threshold to add unsupported gene prediction (bound by 0 and 1)

split_hit=20000 #length for the splitting of hits (expected max intron size for evidence alignments)

min_intron=20 #minimum intron length (used for alignment polishing)

single_exon=1 #consider single exon EST evidence when generating annotations, 1 = yes, 0 = no

single_length=300 #min length required for single exon ESTs if 'single_exon is enabled'

correct_est_fusion=0 #limits use of ESTs in annotation to avoid fusion genes

tries=2 #number of times to try a contig if there is a failure for some reason

clean_try=0 #remove all data from previous run before retrying, 1 = yes, 0 = no

clean_up=0 #removes theVoid directory with individual analysis files, 1 = yes, 0 = no

TMP=/home/sunj/temp #specify a directory other than the system default temporary directory for temporary files
